# Persistent homology analysis distinguishes pathological bone microstructure in non-linear microscopy images

**DOI:** 10.1101/2022.07.19.500658

**Authors:** Ysanne Pritchard, Aikta Sharma, Claire Clarkin, Helen Ogden, Sumeet Mahajan, Rubén J. Sánchez-García

## Abstract

We present a topological method for the detection and quantification of bone microstructure from non-linear microscopy images. Specifically, we analyse second harmonic generation (SHG) and two photon excited autofluorescence (TPaF) images of bone tissue which capture the distribution of matrix (fibrillar collagen) structure and autofluorescent molecules, respectively. Using persistent homology statistics with a signed Euclidean distance transform filtration on binary patches of images, we are able to quantify the number, size, distribution, and crowding of holes within and across samples imaged at the microscale. We apply our methodology to a previously characterized murine model of skeletal pathology whereby vascular endothelial growth factor expression was deleted in osteocalcin-expressing cells (OcnVEGFKO) presenting increased cortical porosity, compared to wild type (WT) littermate controls. We show significant differences in topological statistics between the OcnVEGFKO and WT groups and, when classifying the males, or females respectively, into OcnVEGFKO or WT groups, we obtain high prediction accuracies of 98.7% (74.2%) and 77.8% (65.8%) respectively for SHG (TPaF) images. The persistence statistics that we use are fully interpretable, can highlight regions of abnormality within an image and identify features at different spatial scales.

## 1 Introduction

A wide range of imaging techniques are used to investigate and analyse bone tissue to quantify the effects on structure and morphology as a result of genetic alterations or disease, both for research and diagnostic purposes. Non-linear microscopy imaging techniques such as two photon excitation fluorescence (TPEF) and second harmonic generation (SHG) are well-suited to imaging biological tissue in their native state without the need to label components with dyes or stains to generate image contrast, yet can provide chemical and structural details of samples. These label-free non-linear techniques use a near infrared pulsed laser source to expose samples instantaneously to high-intensity light while keeping the average (total) power within the damage threshold of the tissue. Both TPEF images and SHG images are greyscale, and each image can be encoded as an *n* × *m* matrix with integer pixel intensities between 0 (black) and 255 (white). In TPEF [1, 2], molecules are excited to a higher energy level by near simultaneous absorption of two photons, causing the excited molecules to ‘fluoresce’ spontaneously i.e. emit a single photon, with more energy than one of the absorbed photons. We use TPEF images of the natively fluorescent (autofluorescent) molecules in bone, which we refer to as two photon auto-fluorescence images (TPaF), which provide detail of the overall morphology. In SHG imaging, the second harmonic wave is only generated by media that are non-centrosymmetric i.e. lack structural inversion symmetry in the plane [3], such as crystalline quartz, myosin, tubulin and fibrillar collagen [4, 5]. Therefore, SHG is ideal for selectively imaging collagen fibres and their distribution in human skin, tissues, tendons and bones [4]. Collagen is the most abundant protein in human tissue and is a predominant component of the extracellular matrix in tissues. The role of the extracellular matrix in health conditions and disease processes is increasingly becoming established and is now often studied by using SHG imaging [6, 7]. In bone tissue, SHG specifically images type I collagen fibres and their distribution, that is, the bulk of the bone matrix. Together, two photon autofluorescence (TPaF) and second harmonic generation (SHG) form a set of complementary techniques to image the chemical and structural changes, providing a label-free imaging readout of the bone microstructural properties (characterised by the absence of TPaF and SHG signals).

Correspondingly to these imaging techniques, there is a need for automated and quantitative methods that can effectively analyse the structure in these images in a biologically meaningful way. Topological data analysis (TDA) refers to a collection of topology-based methods that are able to quantify ‘shape’ within data, so are a natural choice to capture structural differences in the samples without requiring large sample sizes and can provide interpretable summaries in the context that each statistic is a measure of specific features in the images (Table 1). This is in contrast to data-driven techniques such as machine learning, which often lack interpretability and require a large training data set for similar classification tasks, something typically infeasible in animal or human research.

**Table 1.** Interpretation of persistence statistics. An advantage of using persistent homology with a SEDT filtration is its interpretability. Here we list the persistent statistics that we use and their morphological interpretation. We use ‘Q.’ for quadrant, ‘p.’ for percentile, ‘st. dev.’ for the standard deviation, IQR for the interquartile range, and micro-hole radius or size refers to the radius of the largest inscribed (open) circle. The inter-micro-hole distance is half the distance from one micro-hole to the next nearest micro-hole.

| $H_*$ | Q | Persistent Statistic | Interpretation |
| --- | --- | --- | --- |
| $H_0$ | 2 | number of points | number of micro-holes |
| $H_0$ | 2 | mean birth | mean micro-hole radius (of largest inscribed circle) |
| $H_0$ | 2 | median birth | median micro-hole radius |
| $H_0$ | 2 | 25p. birth | minimum radius of largest 25% of micro-holes |
| $H_0$ | 2 | IQR birth | range in radii of middle 50% of micro-holes |
| $H_0$ | 2 | st. dev. birth | spread of micro-hole radii |
| $H_0$ | 2 | mean death | average micro-hole crowding (mean of inter-micro-hole distance) |
| $H_0$ | 2 | median death | median inter-micro-hole distance |
| $H_0$ | 2 | 25p. death | maximum of smallest 25% of the inter-micro-hole distances |
| $H_0$ | 2 | IQR death | range in the middle 50% of the inter-micro-hole distances |
| $H_0$ | 2 | st. dev. death | spread of inter-micro-hole distances |
| $H_0$ | 2 | persistent entropy | diversity of micro-hole lifetimes |
| $H_0$ | 2 | number micro-holes $r \geq 2$ | number of micro-holes with radius greater than or equal to 2 |
| $H_0$ | 2 | number micro-holes $r < 2$ | number of micro-holes with radius less than 2 |
| $H_1$ | 1 | number of points | number of bone regions surrounded by micro-holes |
| $H_1$ | 1 | mean birth | mean distance from micro-holes at which loop is formed |
| $H_1$ | 1 | median birth | middle of the distances from micro-holes at which loop is formed |
| $H_1$ | 1 | 25p. birth | distance 25% through distances from micro-holes when loop is formed |
| $H_1$ | 1 | IQR birth | range of middle 50% of distances from micro-holes when loop is formed |
| $H_1$ | 1 | st. dev. birth | spread of distances from micro-holes at which loop is formed |
| $H_1$ | 1 | mean death | mean of the radii of regions surrounded by micro-holes |
| $H_1$ | 1 | median death | radius of middle-sized bone region surrounded by micro-holes |
| $H_1$ | 1 | 25p. death | max radius of smallest 25% of bone regions surrounded by micro-holes |
| $H_1$ | 1 | IQR death | range in radii of middle 50% of bone regions surrounded by micro-holes |
| $H_1$ | 1 | st. dev. death | spread of radii of bone regions surrounded by micro-holes |
| $H_1$ | 1 | persistent entropy | diversity of lifetimes for bone regions surrounded by micro-holes |

We define the term ‘micro-holes’ on three levels: mathematically, they are connected regions of black pixels in a binary image. For microscopy imaging, micro-holes are regions with low signal, that is, pixels that convert to background in the binary image (where the TPaF signal is autofluorescence, the SHG signal is largely collagen in these samples). At the biological level, for the bone samples we analyse, micro-holes encompass a range of structures including osteocyte lacunae and vascular canals (see [7, 8] for the details). Using a powerful and versatile TDA technique called persistent homology, we can analyse the distribution of micro-holes in cortical bone samples i.e. we can quantify the number, size, connectivity, crowdedness and organisation of the micro-holes in the sample.

Persistent homology [9, 10] is a topological method which tracks the evolution of connected pieces, loops and higher-dimensional holes of a ‘continuous shape’, i.e. a topological space [11], represented as a discrete structure such as a simplicial or cubical complex, as a scale parameter varies (Fig. 2b). This gives a persistence diagram of points (Fig. 2c), which is a plot of births (scale parameter value at which the feature appears) against deaths (scale parameter value at which the feature disappears, namely the pieces join or the loops or higher-dimensional ‘holes’ fill). To combine persistent homology with classification algorithms, the format of persistence diagrams needs to be adapted into feature vectors, which can be done through the use of persistence statistics and summary functions [12–15], persistence images [16], or persistence landscapes [17, 18], with the former being our choice as we can tailor specific persistent summaries to bone microstructure measures of interest. TDA methods, including persistent homology, are being increasingly applied to biomedical image data including the cortical thickness of brain from magnetic resonance (MR) images in autistic patients [19]; chronic obstructive pulmonary disease [20]; identifying regions of interest in images of the colon [21]; the classification of diabetic retinopathy images [12]; hepatic lesions [22]; skin lesions [23]; and endoscopy images of the stomach [24]. Previous TDA applications to biomedical images relate to X-ray, Computed Tomography (CT) and MRI techniques. As far as we know, our work is the first to use persistent homology for TPaF and SHG microscopy images of biological samples.

**Figure 1.**
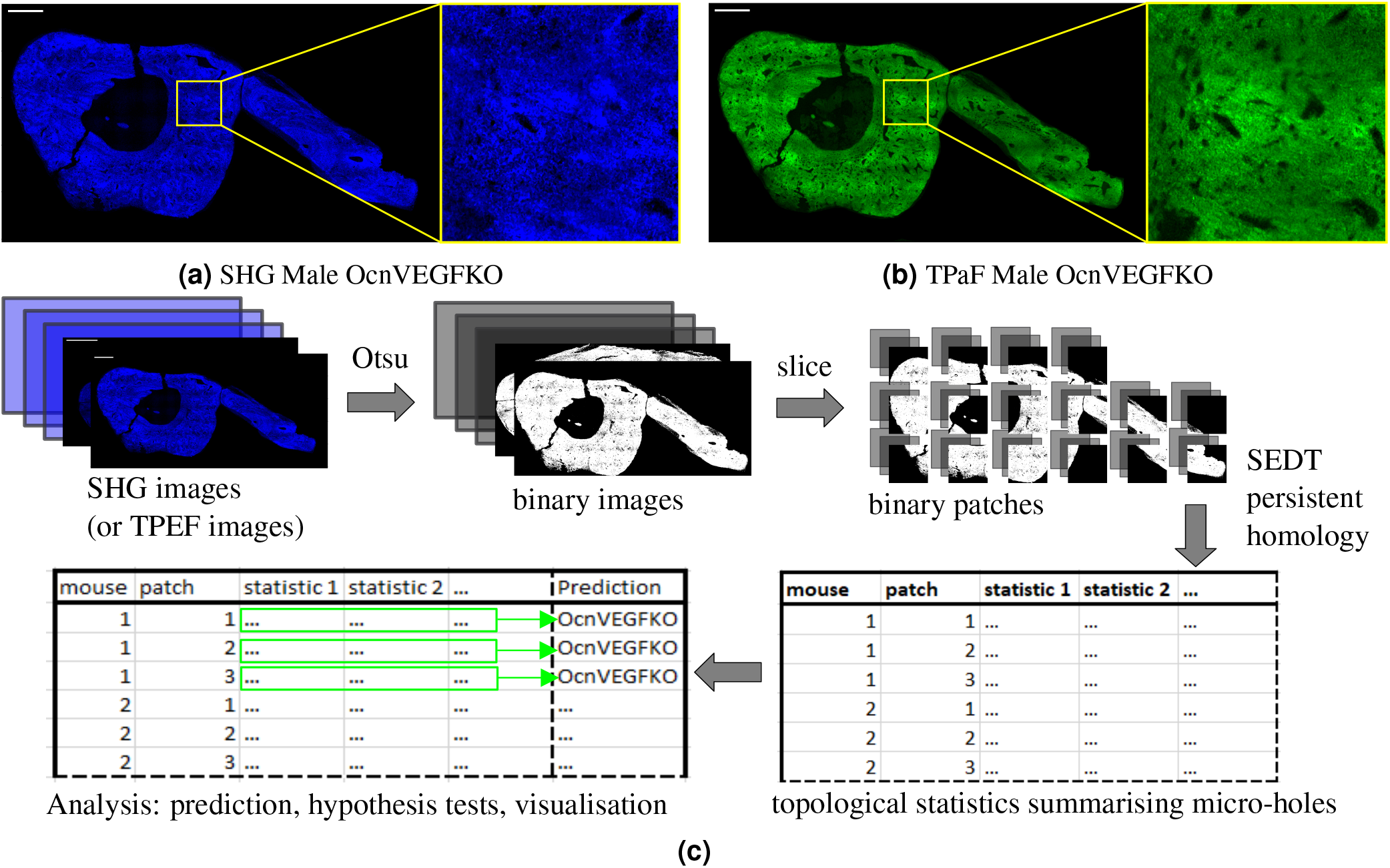
Non-linear microscopy images adapted from [7] and methodology schematic. Tibiofibular junction (TFJ) bone section of a male Osteocalcin-specific *Vegf* knockout (OcnVEGFKO) with SHG (a) and TPaF (b) images from a custom multiphoton microscope [7], with zoomed regions. Scale bars are 250μm. Schematic in (c) gives a high level overview of the method process.

**Figure 2.**
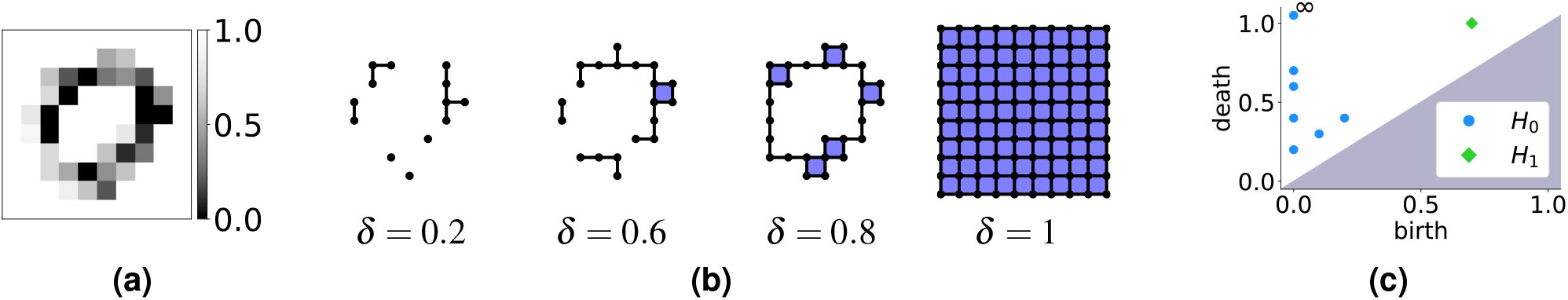
Cubical levelset filtration of a greyscale image. **(a)** Example greyscale image with pixel intensity values between 0 (black) and 1 (white), as in the colour bar. **(b)** Some ordered steps of the cubical levelset filtration of the image in (a). As the threshold parameter *δ* increases from 0 to 1, pixels (of intensity lower than *δ*) are included as points (0-cubes), with two adjacent pixels joined by edges (1-cubes), and 4 adjacent pixels joined by squares (2-cubes). Note the ring we see in the image is captured by the third step of the filtration shown. **(c)** Persistence diagram for *H*_0_ and *H*_1_ of the image (a) with respect to the levelset filtration described in (b). The blue dots (*H*_0_) correspond to connected pieces, which ‘die’ as they join one another. The final connected piece never ‘dies’, so has infinite persistence (marked as ‘∞’). The green diamond (*H*_1_) corresponds to a loop (b, panel 3) capturing the ring structure in the image (a). This loop disappears (‘dies’) as we add more pixels (b, panel 4).

We validate our methodology using previously published TPaF and SHG images of a murine osteocalcin-specific *Vegf* knockout (OcnVEGFKO) bone exhibiting increased cortical porosity [8] in CT images and pathological extracellular matrix organisation [7] (Fig. 1). Our topological analysis (summarised in Figure 4) summarises the micro-holes providing insight into micro-hole organisation and structure (Table 1, Fig. 5). These statistics can be refined to include only features at specific scales of biological interest (Fig. 4c), and can be compared across patches for intra-sample analysis (Fig. 4d). The statistical significance of our topological summaries is tested directly using permutation hypothesis tests, as shown by Table 2. These tests show significant differences in persistence statistics of image patches (Table 2). Finally, we train a support vector classifier to predict whether unseen patches of samples are from the WT (control) or OcnVEGFKO (test) categories. The performance results, shown in Table 3, confirm the strong predictive power of the topological features, particularly from SHG images.

**Figure 3.**
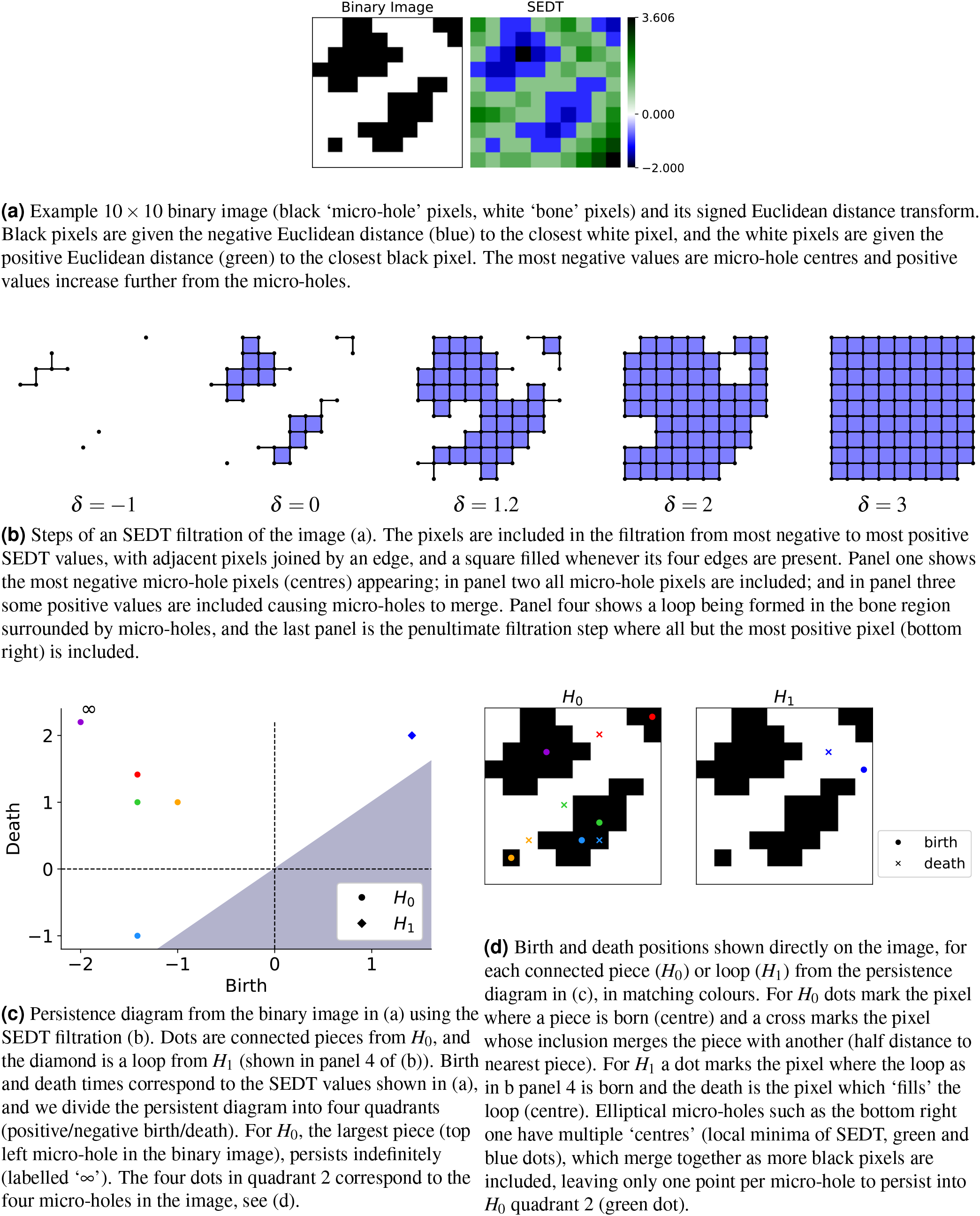
Cubical SEDT filtration of a binary image and persistent homology diagram.

**Figure 4.**
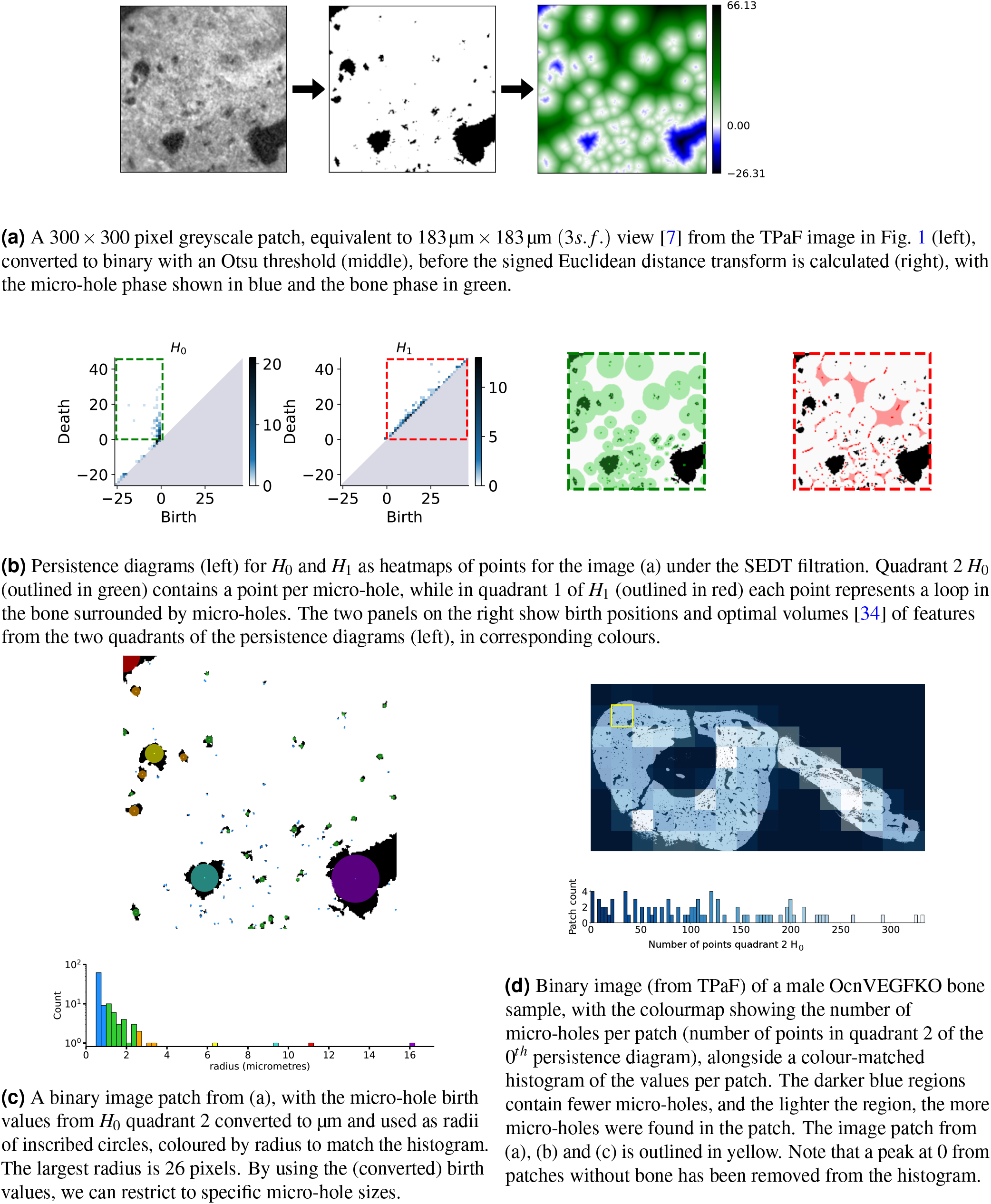
Persistent homology micro-hole structure analysis in a greyscale microscopy image.

**Figure 5.**
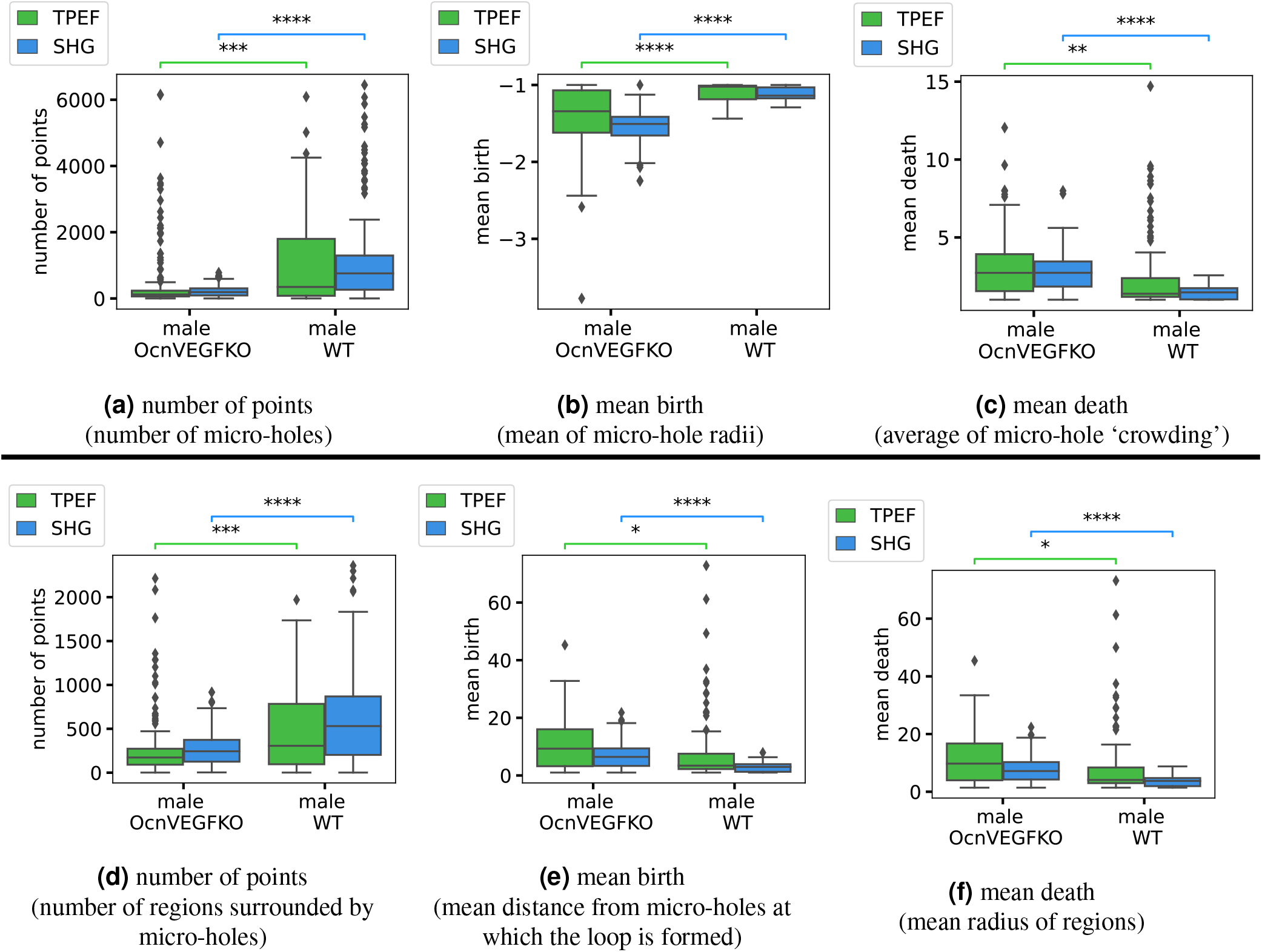
Box plots of persistence statistics per image patch, with three (a-c) from quadrant 2 of *H*_0_ which summarise the micro-holes and three (d-f) persistence statistics from quadrant 1 of *H*_1_, which summarise the loops in the filtration that are regions of bone surrounded by micro-holes. The box plots show the median and quartiles, with outliers marked as points outside of 1.5 times the interquartile range. We annotate their significance using the adjusted *p*-values comparing male OcnVEGFKO to male WT on each imaging type (TPaF or SHG). Here the stars indicate significance levels: ‘ns’ if it is not significant 0.05 < *p* ≤ 1, ‘*’ if 0.01 < *p* ≤ 0.05, ‘**’ if 0.001 < *p* ≤ 0.01, ‘***’ if 0.0001 < *p* ≤ 0.001, and ‘****’ if *p* ≤ 0.0001. We include only the male patches and two of eight significance annotations for readability; for the full *p*-value results, see Table 2.

**Table 2.** Benjamini-Hochberg adjusted pseudo *p*-values (3 d.p.) from permutation tests for 28 test statistics with paired tests to compare the imaging types and tests between OcnVEGFKO (labelled KO above) and WT males and female groups using either the SHG or TPaF images. To see if there is a significant difference from the imaging technique we paired the TPaF and SHG image patches per statistic. Also, for SHG and TPaF separately we tested the hypothesis that the distributions are the same for males between OcnVEGFKO (test) and WT (WT); females between OcnVEGFKO and WT; OcnVEGFKO between male and female; and WT between male and female. All *p*-values in **bold** are significant at the 5% level. The adjustments were made separately for the statistics from TPaF and SHG images in the OcnVEGFKO/WT M/F group comparisons. Here *H*_*_ is the homology group, Q is the quadrant and we have included statistics from the *H*_0_ quadrant 2 (separate micro-holes) and from *H*_1_ quadrant 1 (loops in bone surrounded by micro-holes). We write p. for percentile, *r* for radius and st. dev. is the standard deviation. For the interpretation of each persistent statistic, see Table 1.

| $H_*$ | Q | Persistent Statistic | All<br>SHG<br>v TPaF | Male only:<br>KO v WT | | Female only:<br>KO v WT | | KO only:<br>Male v Female | | WT only:<br>Male v Female | |
| --- | --- | --- | --- | --- | --- | --- | --- | --- | --- | --- | --- |
|  |  |  |  | SHG | TPaF | SHG | TPaF | SHG | TPaF | SHG | TPaF |
| $H_0$ | 2 | number of points | 0.628 | **0.000** | **0.000** | **0.001** | 0.585 | **0.000** | 0.092 | 0.764 | 0.523 |
| $H_0$ | 2 | pers. entropy | **0.015** | **0.000** | **0.001** | 0.528 | 0.243 | **0.000** | **0.009** | 0.138 | 0.586 |
| $H_0$ | 2 | mean birth | **0.000** | **0.000** | **0.000** | **0.000** | 0.104 | **0.000** | **0.000** | **0.009** | 1.000 |
| $H_0$ | 2 | median birth | **0.000** | **0.000** | **0.037** | 1.000 | 1.000 | **0.000** | **0.026** | 1.000 | 1.000 |
| $H_0$ | 2 | 25p. birth | **0.000** | **0.000** | **0.000** | **0.001** | 0.104 | **0.000** | **0.000** | **0.010** | 0.523 |
| $H_0$ | 2 | IQR birth | **0.000** | **0.000** | **0.000** | **0.001** | 0.104 | **0.000** | **0.000** | **0.008** | 0.557 |
| $H_0$ | 2 | st. dev. birth | **0.007** | **0.000** | **0.000** | **0.000** | 0.585 | **0.000** | **0.000** | 0.764 | 0.430 |
| $H_0$ | 2 | mean death | **0.000** | **0.000** | **0.009** | 0.992 | 0.365 | **0.000** | **0.000** | **0.000** | 0.384 |
| $H_0$ | 2 | median death | 0.870 | **0.000** | **0.004** | 0.776 | 0.104 | **0.000** | **0.000** | **0.000** | 0.384 |
| $H_0$ | 2 | 25p. death | **0.000** | **0.000** | 0.135 | 0.528 | 0.104 | **0.000** | 0.091 | 0.058 | 0.384 |
| $H_0$ | 2 | 75p. death | **0.001** | **0.000** | **0.002** | 0.992 | 0.980 | **0.000** | **0.000** | **0.006** | 0.586 |
| $H_0$ | 2 | IQR death | **0.000** | **0.000** | **0.002** | 0.992 | 0.980 | **0.000** | **0.000** | **0.007** | 0.658 |
| $H_0$ | 2 | st. dev. death | **0.000** | **0.000** | **0.033** | 0.992 | 0.104 | **0.000** | **0.019** | **0.000** | 0.384 |
| $H_0$ | 2 | number holes $r \geq 2$ | **0.000** | 0.158 | **0.000** | **0.000** | **0.003** | 0.149 | **0.019** | **0.000** | 0.384 |
| $H_0$ | 2 | number holes $r < 2$ | 0.344 | **0.000** | **0.000** | **0.000** | 0.585 | **0.000** | 0.091 | 0.808 | 0.523 |
| $H_1$ | 1 | number of points | **0.000** | **0.000** | **0.000** | 0.144 | 0.718 | **0.000** | 0.085 | 0.331 | 0.430 |
| $H_1$ | 1 | pers. entropy | **0.000** | **0.000** | **0.001** | 0.995 | 0.585 | **0.000** | **0.019** | 0.197 | 0.982 |
| $H_1$ | 1 | mean birth | **0.000** | **0.000** | **0.037** | 0.797 | 0.392 | **0.000** | **0.019** | **0.000** | 0.430 |
| $H_1$ | 1 | median birth | **0.000** | **0.000** | 0.182 | 0.776 | 0.585 | **0.000** | 0.132 | **0.000** | 0.384 |
| $H_1$ | 1 | 25p. birth | **0.000** | **0.000** | 0.471 | 0.470 | 0.640 | **0.000** | **0.001** | **0.000** | 0.384 |
| $H_1$ | 1 | IQR birth | **0.000** | **0.000** | **0.017** | 0.995 | 0.585 | **0.000** | 0.105 | **0.000** | 0.700 |
| $H_1$ | 1 | st. dev. birth | **0.000** | **0.000** | **0.001** | 0.991 | 0.104 | **0.000** | 0.177 | **0.000** | 0.700 |
| $H_1$ | 1 | mean death | **0.000** | **0.000** | **0.037** | 0.981 | 0.392 | **0.000** | **0.019** | **0.000** | 0.430 |
| $H_1$ | 1 | median death | **0.000** | **0.000** | 0.170 | 0.776 | 0.585 | **0.000** | 0.141 | **0.000** | 0.384 |
| $H_1$ | 1 | 25p. death | **0.000** | **0.000** | 0.478 | 0.797 | 0.640 | **0.000** | **0.002** | **0.001** | 0.384 |
| $H_1$ | 1 | 75p. death | **0.000** | **0.000** | **0.037** | 0.960 | 0.585 | **0.000** | **0.026** | **0.000** | 0.523 |
| $H_1$ | 1 | IQR death | **0.000** | **0.000** | **0.017** | 0.991 | 0.585 | **0.000** | 0.098 | **0.000** | 0.700 |
| $H_1$ | 1 | st. dev. death | **0.000** | **0.000** | **0.001** | 0.992 | 0.104 | **0.000** | 0.166 | **0.000** | 0.700 |

**Table 3.**
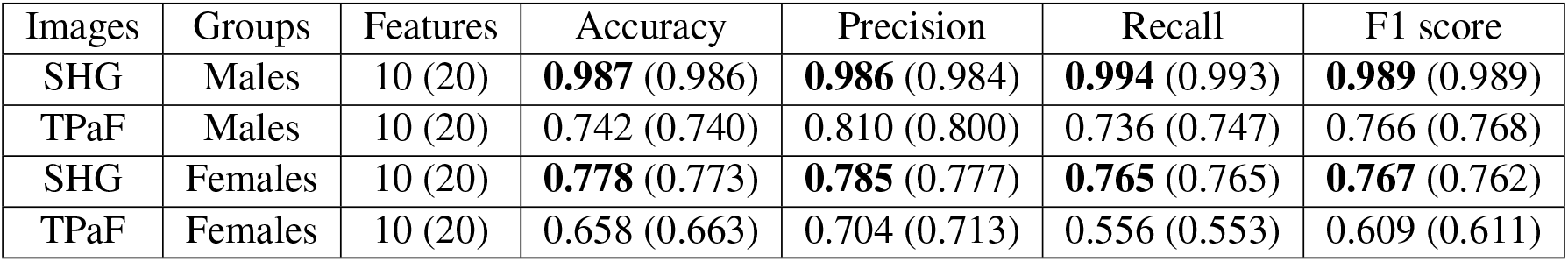
Performance results (for OcnVEGFKO vs WT mice image patches) of an SVC (support vector classifier) with an RBF (radial basis function) kernel (*C* = 3) with stratified 10-fold cross validation, using 10 or 20 persistence features (see main text) to predict whether an image patch in the test set is from a OcnVEGFKO or WT mouse. Classifiers with 10 features use only quadrant 2 of *H*_0_, whereas 20 features adds additional features from quadrant 1 of *H*_1_, with no improvement. All given measures are mean measures over 100 repeats. Numbers in **bold** show the best performance results (SHG images, 10 features).

| Images | Groups | Features | Accuracy | Precision | Recall | F1 score |
| --- | --- | --- | --- | --- | --- | --- |
| SHG | Males | 10 (20) | **0.987** (0.986) | **0.986** (0.984) | **0.994** (0.993) | **0.989** (0.989) |
| TPaF | Males | 10 (20) | 0.742 (0.740) | 0.810 (0.800) | 0.736 (0.747) | 0.766 (0.768) |
| SHG | Females | 10 (20) | **0.778** (0.773) | **0.785** (0.777) | **0.765** (0.765) | **0.767** (0.762) |
| TPaF | Females | 10 (20) | 0.658 (0.663) | 0.704 (0.713) | 0.556 (0.553) | 0.609 (0.611) |

Adapting persistent homology to analyse biologically or clinically relevant features in non-linear microscopy images expands the set of available tools that capture skeletal morphology. The topological approach that we present is not limited to this bone cell specific OcnVEGFKO or mouse models, and can be used in biological sample images where the analysis of micro-holes or hole-like structures may be required, including in a diagnostic context.

Our article is organised as follows. A detailed description of our methodology is given in section 2, including the specimen preparation, SHG and TPaF imaging, pre-processing of the images, a short introduction to persistent homology, a description of the filtration called the signed Euclidean distance transform (SEDT) that we use as input for the persistent homology, the persistence statistics per quadrant, the permutation hypothesis tests, and the classification using a support vector machine. In section 3, we present the results of our topological methodology applied to mice bone images, including the outcome of the hypothesis tests, the interpretation of the most meaningful persistence statistics between groups of mice, and the results of the classification. The article finishes with a discussion and some conclusions in sections 4 and 5.

## 2 Methods

### 2.1 Specimen Preparation

The data presented herein were acquired previously on samples used in the studies by Goring et al and Sharma et al [7, 8]. The tibiae were derived from 16 week-old littermate matched mice evenly split between male and female, and OcnVEGFKO and WT [7, 8]. Briefly [8], the tibiae were fixed in 4% paraformaldehyde for 48 hours, dehydrated in 70% ethanol and embedded in poly-methyl methacrylate, prior to taking 5μm thick sections from the tibiofibular junction (TFJ). The use of animal tissue was carried out in compliance with the Animals (Scientific Procedures) Act 1986 and the regulations set by the UK home office, as in [7, 8].

### 2.2 SHG and TPaF Microscopy Imaging

Two photon excitation autofluorescence (TPaF) and (circularly polarised) second harmonic generation (SHG) images of each bone section were previously acquired by Sharma et al [7] using a custom multiphoton microscopy setup. For full details of the imaging system and acquisition methodology see [7]. Example images are shown in Fig. 1. The images are 2D greyscale images with integer pixel intensities between 0 (black) and 255 (white). The TPaF images show the mineralised matrix and the SHG images capture the collagen fibres. Per imaging technique, there are 12 images of the 2-3mm bone sections described in section 2.1. Note that the breaks in the bone are irrelevant for our microstructure analysis.

### 2.3 Preprocessing Images

We thresholded all images of the mouse samples to binary using Otsu’s method [25], which divides the image pixels into two classes (black and white) by maximising the between-class variance. We trimmed the binary images of empty borders, padded them to an integer multiple of the desired patch shape, and split each image into 300 × 300 pixel patches, which is equivalent to a 183μm × 183μm (3*s. f*.) view [7]. Note the number of patches is not an input but is determined by patch size and how many fully background patches are discarded. For these images, this scale allows for the largest meaningful features to be present in a patch. The patches were taken with a stride of 300 pixels to ensure they did not overlap, and we discarded any patches that were entirely background. This gave us a total of 475 patches and 476 patches from the original 12 TPaF and SHG images respectively, which were treated as separate sets of images in the analysis process. Note that edge patches have varying amounts of bone and, due to the irregular sample shape and to differences in the size of the bone sections, each individual image may produce a different number of patches. For the SHG (TPaF) images respectively there were 93 (93) female KO patches, 103 (101) female WT patches, 163 (160) male KO patches, and 117 (121) male WT patches.

### 2.4 Topological Data Analysis Method

Persistent homology tracks the evolution of connected pieces, loops and higher-dimensional ‘holes’ by calculating the homology groups of an increasing sequence of complexes (called a filtration) built using a scale parameter (Fig. 2b). We used cubical homology (section 2.4.1) and a signed Euclidean distance transform (SEDT) filtration (Fig. 3, section 2.4.2) on pre-processed binary image patches (Fig. 4a, section 2.3). The quadrants of the resulting persistence diagrams (Fig. 4b) quantify specific features, such as separate micro-holes, connected networks of micro-holes, and loops in bones surrounded by micro-holes (Fig. 4b, Table 1). Using this, we calculated summary statistics per patch (Fig. 5, section 2.4.3), which quantify the quantity, size and organisation of the micro-holes within each patch, and can be visualised on the whole sample to highlight regions of interest (Fig. 4d).

All calculations were done in python 3 on a laptop with 5 cores and 8GB RAM running Windows 10. Image data and dataframes were handled in PIL, numpy, and pandas, persistent homology was calculated using the homcloud package, and for classification we used sklearn. The figures were created using matplotlib, seaborn and statannot. All computer code, documentation and an example computation are available on github [26]. Calculating the persistent homology, the longest part of the computation, took under 2 seconds per patch, and can be parallelised.

#### 2.4.1 Persistent homology of cubical complexes

The first step to apply a topological method to a data set is to build a ‘shape’ (formally, a topological space) from the data points and some measure of similarity or distance between them. We build a topological space by gluing together basic building blocks (typically triangles or squares, and their higher-dimensional counterparts) with the data points as vertices. For images (pixels on a rectangular grid), using squares, cubes, etc., is the most natural option, resulting in what we call a cubical complex. More precisely, a cubical complex [27] is a topological space made of any number of *n*-cubes (0-cubes are points, 1-cubes are edges, 2-cubes are squares, 3-cubes are cubes, etc.), for different *n*, glued together along their faces (the vertices of an edge, the sides of a square or a cube, etc.) and embedded in ℝ^d^ (for *d* large enough). A cubical complex is a combinatorial model of a topological space, or, in our case, an image. See Fig. 2b for some examples, and [27] for a formal definition.

Suppose that we want to encode a greyscale image as a cubical complex, taking into account both the pixel positions and their intensities (0 black to 255 white). A standard way is using a levelset filtration: we only include pixels below a certain threshold intensity level, and always join adjacent pixels (pixels whose coordinates differ by exactly 1), filling in squares (or cubes, for 3-dimensional images) when all the edges are present. As we increase the threshold intensity level, we include more and more pixels and obtain a sequence of increasingly larger complexes, called a filtration (see Fig. 2b for an example). There are other ways to construct a filtration of cubical complexes from an image and we use the Signed Euclidean Distance Transform (SEDT) filtration, explained in section 2.4.2.

Before describing the SEDT filtration, we will briefly and informally discuss persistent homology (for a formal treatment, see e.g. [9, 28, 29]). The homology groups *H_k_*(*X*) of a topological space *X* (such as a cubical complex) detect the number of *k*-dimensional ‘holes’ in *X*, for *k* = 0,1,2,… For *k* = 0, *H*_0_(*X*) detects the connected ‘pieces’ that make up *X*, while *H*_1_ (*X*) detects the cycles (loops) in *X* (see Fig. 2b). The rank of the homology groups (the number of *k*-dimensional ‘holes’) are called the Betti numbers, written *β_k_*, and uniquely determine *H_k_*(*X*) for each *k* (assuming coefficients in a field *F*, which is *F* = ℝ in our case). For full details on the homology of cubical complexes, see [27]. Persistent homology extends homology from a single topological space to a filtration (a nested, increasing sequence of topological spaces). It calculates the homology groups across the filtration, keeping track of when a feature is born (appears) or dies (disappears). The result is encoded in a persistence diagram, such as the one shown in Fig. 2c. Note that, since we analyse 2D images, we only consider homology for dimensions *k* = 0, 1, as the homology is zero for larger *k*. Each persistent diagram is a plot of pairs of points (*b_i_*, *d_i_*) where *b_i_*, *d_i_* are respectively the birth and death time, of the *i^th^ k*-dimensional feature in *H_k_*. A feature can only die after it is born, hence *b_i_* < *d_i_* and all points shown in a persistent diagram must be above the diagonal. It can be shown that all features eventually die (at most when the connectivity between data points is maximal), except in *H*_0_ where a final connected ‘piece’ of *X* remain indefinitely (shown with the symbol ∞ in our persistent diagrams). In Figure 2, we show a sample greyscale image, levelset filtration, and resulting persistent homology diagram, for illustration.

#### 2.4.2 Signed Euclidean distance transform

We analyse the micro-holes in the SHG and TPaF images (which we refer to as ‘microstructure’) of bone samples using persistent homology with a signed Euclidean distance transform (SEDT) filtration (Fig. 3, 4). The SEDT filtration allows us to encode the different structural features of an image as points in specific quadrants of the resulting persistence diagrams. This filtration has been successfully used to study the structure of bead packing, sand packing, limestone and sandstone samples on micro CT images in [30], and to analyse the flow of fluids and trapping of bubbles in sandstone samples in 3D X-ray CT scans [31]. In medical imaging, the SEDT filtration has been recently adapted to tumour images by Moon et al [32] to study the relation between topological shape features in tumour progression and survival risks. Further, a similar (Manhattan) distance transform filtration approach has been used in [33] to predict transepidermal water loss from skin images.

The input of the signed Euclidean distance transform (SEDT) is a binary (black/white) image, rather than a greyscale image. It takes the binary image and assigns to every pixel in both phases (white, or black, pixels) the shortest Euclidean distance to the opposite phase of the image. Pixels in the white (‘bone’) phase are given a positive distance, and pixels in the black (‘micro-hole’) phase are given a negative distance, as in Fig. 3a. We use persistent homology with a signed Euclidean distance transform filtration as in [31], that is, the filtration where we include pixels from the most negative distance to the most positive distance, with respect to the SEDT. The pixels with the most negative distance values are the micro-hole centres and are included first in the filtration, while the pixels with the greatest positive distances are bone pixels furthest from any micro-holes and are included last in the filtration. In Fig. 3, we show an example of the persistent homology of a binary image with respect to the SEDT filtration in detail. In Fig. 4, we explain the process taking a patch of a greyscale image to binary, before calculating the persistent homology using an SEDT filtration, and showing the quadrants of the persistence diagrams and how the features captured in each quadrant relate back to the micro-holes in the binary image.

Let us discuss a few properties of the SEDT persistent homology which are relevant to our analysis (cf. section 2.4.3 and Table 1). We can divide the persistent diagram into four quadrants (positive/negative birth/death), which we label 1 to 4 from the top right, proceeding anti-clockwise (see Fig. 3c). Since all points are above the diagonal, there are no points in quadrant 4. All black pixels (and thus all micro-holes, which we define formally as connected regions of black pixels in the micro scale binary images of the samples) are born by *δ* = 0 (let us call *δ* the parameter in the SEDT filtration), hence there are no *H*_0_ points in quadrant 1. There can be *H*_0_ points in quadrant 3, for connected micro-holes born at different pixels (see Fig. 3c) and merged later (at most by *δ* = 0, when all black (micro-hole) pixels are included). More irregular (e.g. less ‘circular’) micro-holes can be born at different pixels (at each local minimum of the SEDT distance within the same micro-hole). By *δ* = 0, all micro-holes have been included: the number of *H*_0_ points in quadrant 2 (including the point labelled ∞) equals the number of micro-holes, and each of the remaining points in quadrant 3 corresponds to a unique point (micro-hole) in quadrant 2. Moreover, the birth of each of these points equals the radius of the largest (open) circle that can be inscribed in the micro-hole, which follows from the definition of the SEDT. We refer to this quantity as the *size* of the micro-hole. Hence, the larger the micro-hole, the earlier it is born (more negative *δ*), with the largest micro-hole appearing first and persisting indefinitely in the filtration (labelled ‘∞’ in the persistent diagram). Although not entirely obvious, the introduction of white (bone) pixels cannot create new connected pieces: a new white pixel will join its closest micro-hole(s), possibly merging two or more micro-holes. All in all, the persistence diagram for the 0^*th*^ homology group contains the disconnected micro-holes in quadrant 2 and the connected micro-holes in quadrant 3, with empty quadrants 1 and 4. Finally, note that the death value for micro-holes in quadrant 2 of *H*_0_ is half of the shortest distance from the micro-hole to its nearest micro-hole (the distance between two micro-holes *A* and *B* is the minimum distance between a pixel in *A* and a pixel in *B*), as both micro-holes expand by including the white pixels between them and thus it is a measure of micro-hole ‘crowdedness’. To visualise the features from the quadrants of the persistence diagrams under the SEDT filtration, see Fig. 4b. The birth pixels and optimal volumes [34] of the features have been plotted on the images for *H*_0_ quadrant 2 (green), and *H*_1_ quadrant 1 (red), using homcloud.

For the 1^*st*^ homology group (*H*_1_), any loop born before *δ* = 0 would be a loop of black (micro-hole) pixels, containing white (bone) pixels, so it will not die until some *δ* > 0. Therefore, there are no *H*_1_ points in quadrant 3, but there could be points in quadrant 2 (micro-hole loops). Points in quadrant 1 of *H*_1_ correspond to loops in the bone between micro-holes, that is, white (bone) pixels connecting black regions (micro-holes) in a ring (loop) fashion (see Fig. 3), with larger loops indicating regions with sparser micro-hole distribution.

For reproducibility, we include the following technical note. The 0^*th*^ persistence diagram contains a point with the most negative birth value (known as ‘essential birth’) and infinite death (labelled ‘∞’ in Fig. 3c), which naturally falls in quadrant 2 and corresponds to the largest micro-hole (connected region of black pixels) in the binary image patch. These points were omitted when calculating the statistics, since for most patches, the largest negative distance was a point in the background region surrounding the bone sample.

#### 2.4.3 Persistence statistics

For each image patch, we calculate the persistence statistics for quadrant 2 of the 0^*th*^ persistence diagram, and for quadrant 1 of the 1^*st*^ persistence diagram, as these are the most informative quadrants for our analysis (see discussion in section 2.4.2). Namely, for each quadrant of interest, we considered the following summary statistics, calculated per image patch, per homological dimension (*H*_0_ and *H*_1_), and for both birth and death distributions: mean, median, quartiles, interquartile range (IQR) and standard deviation. We also calculated, for each image patch and dimension, the number of points *n* in the persistent diagram, the total persistence 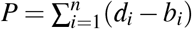, and the persistent entropy [35, 36] 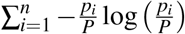. The latter is the Shannon entropy [37] of the persistent values *d_i_* — *b_i_* as a probability distribution (divided by the total persistence *P*), and measures the ‘diversity’ of the persistence [23], that is, how close is the uniform distribution (where all features are equally persistent).

Each of the chosen persistent statistics has a direct interpretation in terms of the micro-hole (bone) structure, as shown in Table 1. Moreover, we can restrict our attention to features within a specific scale of interest, for example micro-holes of a specific size (see Fig. 4c). Further, by focusing on a persistent statistic of interest, we can give image patches a score, highlighting regions of interest within the sample (see Fig. 4d).

### 2.5 Hypothesis Tests

We determine the statistical significance of the persistence statistics described in section 2.4.3 in our data set of image patches using permutation hypothesis tests [38]. Namely, we test the null hypothesis that two groups (e.g. female test (OcnVEGFKO) v. female control (WT)) have identical distributions for the given persistence statistics (e.g. mean birth in *H*_0_ quadrant 1). In the following sections, OcnVEGFKO is referred to as ‘test’ and WT as ‘control’, as the method is independent of the mouse model. The result is a pseudo *p*-value, which is the probability that the difference in the mean statistic between the two shuffled groups is at least as large as the difference we observe, if the null hypothesis that the two groups have the same distribution is true. If this probability is very small, we reject the hypothesis that the distributions are the same for the two groups.

To calculate the pseudo p-value, we randomly assign image patches to two groups (of the same size as the two groups we are comparing) and calculate the difference between the means of the given statistic for the two randomly selected groups. By repeating this step a large number of times (*n* = 10,000 in our tests), we obtain a distribution of differences in means. The pseudo *p*-value is the proportion of the shuffled differences in means which fall above the observed difference in our initial groups. That is, the probability of having observed that difference between two randomly selected groups under the hypothesis that the two groups have identical distributions. This is an estimate of the *p*-value (thus called pseudo *p*-value), since computing all possible permutations is infeasible. In particular, a pseudo *p*-value of zero should be interpreted as a very small non-zero value.

We have also included paired permutation hypothesis tests for each of the 26 statistics to determine if there are significant differences between the statistic calculated on the TPEF and SHG images. We test the null hypothesis that the statistic has identical distributions from the TPEF and SHG imaging types. Each statistic is paired per image patch per sample and are shuffled (*n* = 10,000 times) between the TPEF and SHG statistic column. For each shuffle we calculate the mean of the shuffled differences normalised by the standard deviation of the shuffled differences. We use a two-tailed test and the *p*-value is the proportion of shuffled differences more extreme than the measured difference.

When comparing a large number of persistence statistics using hypothesis tests, we should account for the multiple comparison problem. Given that we do not intend to claim biological discoveries based on a single statistic, the effect that a single false positive has for our experiment is not overly damaging. Thus, for this data, we chose a less-strict measure by using a Benjamini-Hochberg *p*-value adjustment [39] which controls the false discovery rate (expected proportion of all positives that are false) 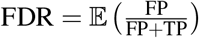, where TP and FP are the number of true positives and false positives respectively.

### 2.6 Classification

In addition to the hypothesis tests (section 2.5), we assessed the usefulness of the persistent statistics for classification purposes by training a classifier on the images to predict whether unseen patches are from our test (OcnVEGFKO) or control (WT) set. That is, we addressed the question whether all or some of the persistent statistics that we considered are enough to distinguish the test condition images from the control images.

We used a support vector classifier (SVC) [40] with a radial basis function (RBF) kernel, which performed better than the more standard linear kernel for our dataset of image patches. Cross-validation [41] was used to split the data into training and test sets, specifically a 10-fold cross-validation stratified into groups where both the training and test sets contain a number of patches from each of the four categories (male OcnVEGFKO, male WT, female OcnVEGFKO, female WT). The resulting prediction accuracy, precision, recall and F1 score [42] on the unseen test sets are given in Table 3.

## 3 Results

Separately for each image type (SHG and TPaF), we took 476 patches from the 12 images, and using hypothesis testing, we compared the persistent statistics (section 2.4.3) for the four groups: male OcnVEGFKO, male WT, female OcnVEGFKO, female WT. Note that we divided the samples not only into WT (control) and OcnVEGFKO (test), but also by sex, due to the known differences per sex of this condition on bone [7, 8]. In this section, we present the results of the hypothesis tests (Table 2), which show where the persistent statistics are significantly different between groups of mice, interpret these differences in the persistence statistics (Fig. 5), and show the performance results of the support vector classifiers (Table 3).

As discussed (section 2.4.2), each micro-hole is represented by a (birth, death) point in quadrant 2 of the *H*_0_ persistent diagram, with the birth the largest distance from the bone matrix (the maximal radius of a circle inscribed in the micro-hole) and the death is (half) the distance at which it merges into other pieces originating from surrounding micro-holes (so the death tells us about the local crowding behaviour). Hence, as in Table 1, the following 13 statistics that summarise this quadrant can summarise the microstructure of the holes: number of points (number of micro-holes), persistent entropy (diversity of persistence), mean and median birth (averages of micro-hole radius), mean and median death (averages of micro-hole crowding), 25^*th*^ percentile of births (minimum radius of largest 25% of micro-holes), 25^*th*^ and 75^*th*^ percentiles of death distribution (crowding bounds of smallest 25% and largest 25% of micro-holes), the interquartile range of births and deaths (radius/crowding difference for middle 50% of micro-holes) and the standard deviation of births/deaths (spread of radius and crowding). Analogous persistent statistics for quadrant 1 of *H*_1_, which captures loops in the bone matrix surrounded by micro-holes, quantify the number and size of uninterrupted regions surrounded by micro-holes. Finally, to show the number of features at a given scale, we also include the number of micro-holes of radius less than 2, and the number of micro-holes of radius greater than or equal to 2, calculated using the births from quadrant 2 of *H*_0_.

In Fig. 5, we show three example persistent statistics for *H*_0_ and three persistent statistics for *H*_1_, which allow us to quantify the differences between the male OcnVEGFKO and WT groups with respect to specific interpretable topological (persistent) features. For the full significant differences per statistic see 2 and for plots of other statistics and the female results please see the supplemental information. The statistics shown are divided into male OcnVEGFKO and WT groups, so there are patches from bone tissue samples from 3 mice, each represented by its respective box plot. Firstly, in Fig. 5a, the male OcnVEGFKO mice have fewer points, hence fewer micro-holes, but the micro-holes are much larger as they have more negative birth times in Fig. 5b, i.e. the micro-hole centres are farther from the bone pixels. The top 25% of micro-holes have a radius larger than the 25^*th*^ percentile for births and the male OcnVEGFKO set have consistently larger radii across the patches. Similarly, the interquartile range (IQR) of births shows the middle 50% of micro-hole sizes have a much greater range of sizes in the OcnVEGFKO male patches compared to the other groups whose separate micro-holes are much more consistent in size. There is a decrease in persistent entropy for the OcnVEGFKO males, which indicates that features are more diverse, as persistent entropy increases with homogeneity of persistence. Importantly, the OcnVEGFKO male patches are also less crowded, which is shown by the increase in mean death time in Fig. 5c, as micro-holes persist to greater distances from the micro-hole before colliding. This is further supported by OcnVEGFKO male patches showing significant increases for deaths in the median, standard deviation, IQR and the 75^*th*^ percentile. It should be noted that, for the SHG images only, there are a few significant differences between female OcnVEGFKO and WT patches, including the number of points, mean birth, IQR, standard deviation and 25^*th*^ percentile for birth.

In addition, for quadrant 1 of the 1^*st*^ persistence diagram (Fig. 5 (d-f)), the male OcnVEGFKO patches contain fewer loops formed by bone matrix regions around micro-holes, as there is a lower number of points (Fig. 5d), which fits with the expectation given there are fewer micro-holes. These loops have a higher mean birth (Fig. 5e), so are born at greater distance from the micro-holes and similarly an increased mean death (Fig. 5f) means they are filled in at higher distance values, so are larger. Overall, this shows that the micro-holes in the OcnVEGFKO male patches are less crowded, as the bone matrix areas between micro-holes are fewer, larger and less interrupted by smaller features.

We show the Benjamini-Hochberg adjusted *p*-values for the permutation tests for our choice of persistence statistics in Table 2. For each imaging type (SHG, and TPaF), we test the hypothesis that the distributions are the same for: males between OcnVEGFKO and WT; females between OcnVEGFKO and WT; OcnVEGFKO between male and female; and WT between male and female. The *p*-values show many significant differences between the male OcnVEGFKO and WT image patches, and between the OcnVEGFKO male and female patches. However, between the female OcnVEGFKO and WT patches, the statistics show only 5 significant differences for the SHG (and none for TPaF). The vast majority of statistics are also significantly different between the OcnVEGFKO male and female patches, both for the SHG and TPaF images. For the SHG images, there are also some significant differences between the male and female WT patches that are not present in the TPaF images, consistent with what we observe in Fig. 5. The SHG images capture the collagen structure of the samples, so the stronger significant differences and improved classification performance shown in Table 3 indicate that the differences are due to the collagen distribution in the bone matrix (see discussion in section 4).

In Table 2 above, the first p-value column gives the results of paired permutation tests, comparing the difference between an SHG patch and the corresponding TPaF patch, for each statistic. The majority of these p-values are significant at the 5% level, demonstrating that the choice of imaging technique results in significantly different persistent statistics. In these paired tests, the statistics relating to the sizes of the micro-holes are significantly different between the imaging techniques. This is to be expected, as the SHG images are detecting micro-holes in the collagen structure, in comparison to micro-holes in the autofluorescence of the TPaF image patches. However, the number of micro-holes was not found to be significantly different between the imaging techniques per patch, indicating a more consistent count of micro-holes between TPaF and SHG images.

By training a classifier on the persistence statistics, we can predict whether an unseen image patch is typical of a condition. Here, we compare image patches from OcnVEGFKO (test) mice to WT (control) mice, per sex. We used a support vector classifier (section 2.6) with 10 of the persistence statistics, per quadrant: number of points, persistent entropy, and the mean, median, standard deviation of births and deaths as well as the 25^*th*^ percentile of births and the 75^*th*^ percentile of deaths. We omit the number of micro-holes with specific radii as they sum to the number of micro-holes, and similarly we also omit the interquartile ranges (IQR) as they are the difference between the quartiles. In addition, the 25^*th*^ percentile of deaths is omitted due to fewer significant differences (see Table 2). The results (Table 3) show a good separation between OcnVEGFKO and WT samples in the males, and reasonably strong results for females, given that there are far fewer significant differences. Further, when including features from quadrant 1 of *H*_1_, results shown in brackets, there is no improvement, which suggests that the features from quadrant 2 of *H*_0_ contain enough topological information on their own.

## 4 Discussion

We have presented a topology-based workflow to quantify morphological features, namely microstructure of the holes in the bone, in non-linear microscopy images. We extracted fully interpretable topological statistics from each image patch (Table 1), which quantify biologically relevant characteristics such as the number, radii and crowding of micro-holes throughout the samples. We demonstrated our methodology on mice bone samples and found significant differences in topological statistics between samples from OcnVEGFKO (test) and WT (control) images (Table 2), in line with previous observations [8]. Furthermore, we were able to accurately classify unseen image patches, using a support vector machine trained with a subset of the topological statistics. On this data, we see improved classification performance on patches from males in comparison to females (Table 3), stemming from the stronger significant differences between male OcnVEGFKO and WT (Table 2). The predictions for the SHG images are notably more accurate, which demonstrates the benefits of using a specialised microscopy technique, such as SHG, which captures the collagen structure.

The main strengths of our methodology are its interpretability, objectivity, and flexibility. Indeed, we were able to directly relate observable, biologically relevant micro-hole characteristics to specific topological statistics and determine whether differences were statistically significant. Namely, using an SEDT filtration, we encoded geometric information, such as radius or inter-micro-hole distances, using topological summaries of persistent homology. Our workflow is automated and objective, relying only on the choice of patch size, which reduces the potential for human error or bias. The persistent homology methodology itself is highly flexible, and can be customised to analyse other morphological characteristics of interest, by tailoring the filtration and choice of persistent statistics, and it can also be used to separate and summarise features of specific sizes (see Fig. 4c or Table 2).

By taking patches of the images, we can analyse the localised microstructure of the samples (such as the average minimal radius of micro-holes in region), at the cost of ignoring the macrostructure (the overall ‘shape’ of the sample), which is irrelevant to our structural analysis. (This is not a real limitation, as our topological workflow can be applied to whole sample images, and the topological summaries adapted to global morphological features of interest.) By using image patches, we also increase the number of images, and, together with the engineered topological features, we make classification tasks feasible on small sample sizes, as demonstrated in our mice data set (Table 3). Moreover, the patch-based approach allows for clear, intuitive visualisations of the topological statistics on the (binary) whole sample image. These visualisations can be used to indicate atypical or interesting regions within a single sample, and allow direct comparison of full sample images across a set of samples, which can highlight consistent regional effects or anomalous samples. Also, it should be noted that the background outside of the sample will register as a single micro-hole per patch containing background, which we mitigated by trimming images to reduce the size of the background component.

There are several existing methods to quantify porosity in bone tissue, including the method in [8, 43] applied to SR CT images of the same samples in [8]. Porosity is a volumetric measure of how porous a sample is. Despite analysing micro-holes in 2D images of 3D ‘pore’ structures, the topological method on SHG images that we have presented is comparable to this ‘geometric’ method on SR CT image data. However, our topological method offers a distinct advantage over existing methods in that it is automated, and encodes a more diverse summary of different microstructure characteristics, including micro-hole crowdedness, which can be used in assessing a wider range of morphological characteristics in bone samples.

An important limitation when using persistent homology is that it only uses one scale parameter (the SEDT values in our analysis), as multi-parameter persistent homology is problematic [44]. In particular, the original pixel intensities are not used beyond thresholding into a binary image. This is necessary for the SEDT filtration, which allows us to separate (and thus summarise) the individual micro-holes (black pixels). Our method, therefore, relies on a good separation of the images to a binary format of bone matrix and micro-hole phase. Note that the existing geometrical method we mentioned above also relies on converting greyscale images into binary.

Finally, in terms of computational cost, the calculation of the persistent statistics took a couple seconds per image patch, and can be run in parallel.

## 5 Conclusion

We have provided a new method for the detection and quantification of micro-hole morphology on microscopy images. Its input is any greyscale image, which is converted into a binary image, and a patch size, and it uses a topological method to summarise micro-holes (regions without signal, or connected pieces of black pixels in the binary image) morphology per image patch. Namely, it detects individual micro-holes (*H*_0_ points) and their radius (largest inscribed circle, *H*_0_ *birth*), as well as the distance to the closest micro-hole (*H*_0_ *death*), in an automatic way. Additionally, we can also summarise the signal region morphology, by detecting signal regions surrounded by microholes (*H*_1_ points), and the formation of loops around signal regions (*H*_1_ *birth*), and the radius of such regions (*H*_1_ *death*). Our topological method is automatic, requiring only a greyscale image and patch size, robust, as persistent homology is known to be robust against noise (a small perturbation of the input results in a small perturbation of the output [45]), and interpretable, in the sense that each topological summary is linked to a micro-hole morphological characteristic (Table 1).

We demonstrate how micro-hole morphology analysis using our topological method is sufficient to detect significant differences, and classify with high predictive power, on transgenic mice bone TPaF and SHG images, despite a small sample size. The use of SHG images provided us with a detailed analysis of micro-holes in the collagen structure, which can be directly compared to those in the autofluorescence of TPaF images. There is a clear benefit to analysing the SHG images, when we look at the comparative strength of the results between the SHG and TPEF images. SHG images are typically used to study collagen structure [46], mostly focusing on analysing or extracting fibres. It would be interesting to combine our topological methods with existing fibre analysis in the context of collagen structure in SHG images, but this is beyond the scope of the present article.

Our topological method provides a new tool in bone image analysis, and is applicable to other samples, materials and imaging techniques where we want to quantify ‘micro-hole’ (lack of signal regions) morphology. We have provided ready-to-use python code for the entire workflow. Note that our topological summaries can be combined with other image analysis features, in particular to improve the predictive performance of a classifier. In that sense, our method can also be seen as high-quality morphological feature engineering for image analysis.

Our results demonstrate the usefulness, versatility, and potential of topological analysis methods in non-linear microscopy imaging. By validating this method on bone samples, we broaden the bone analysis toolkit by adding a flexible topological workflow, and new microstructure topological features for bone. We expect our, and similar, topological methods to be useful in other biological research settings on tissue samples beyond bone, and for a wide range of biological applications. In the future, we plan to adapt our topological method to analyse other microscale morphological characteristics beyond the structure of micro-holes in the image data, by modifying the filtration and customising the topological summaries. One important avenue for future work is to explore the potential for use as a diagnostic aid, and tailor this method to specific diseases which alter bone structure at the microscale within human samples.

## Data Availability

The datasets used and analysed during the current study are available in the figshare repository, under the DOI: https://doi.org/10.6084/m9.figshare.20765659.v1.

## Acknowledgements

YP acknowledges funding from the School of Mathematical Sciences, University of Southampton and EPSRC grant EP/N509747/1. SM acknowledges EPSRC Transformative Healthcare 2050 grant EP/T020997/1. RSG acknowledges support from The Alan Turing Institute under the EPSRC grant EP/N510129/1.

## Author contributions statement

A.S. and C.C. sourced, prepared and imaged the samples, Y.P., R.S.G., H.O. and S.M. conceived the project, R.S.G., H.O. and S.M. supervised the project, Y.P. completed the calculations and prepared the first draft of the manuscript. All authors reviewed and contributed to the manuscript.

