## Supplementary Information for "Persistent homology analysis distinguishes pathological bone microstructure in non-linear microscopy images"

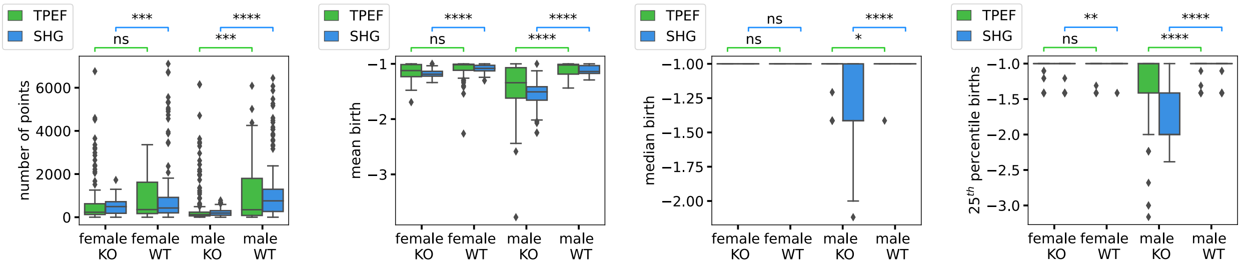


**(a)** number of points **(b)** mean birth **(c)** median birth **(d)** $25^{th}$ percentile birth

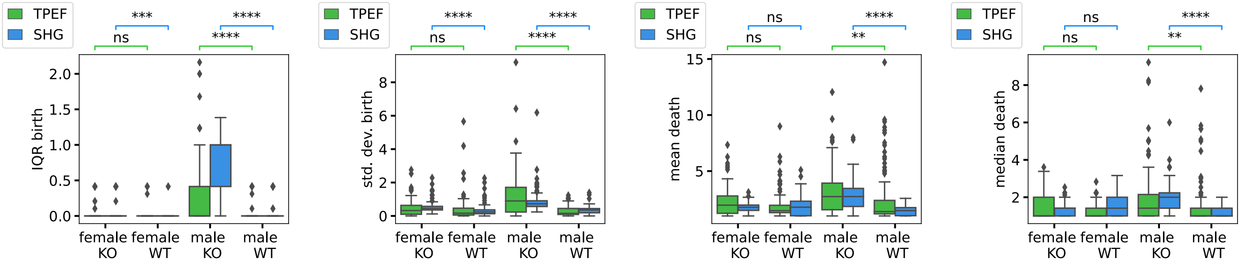


**(e)** IQR birth **(f)** std. dev. birth **(g)** mean death **(h)** median death

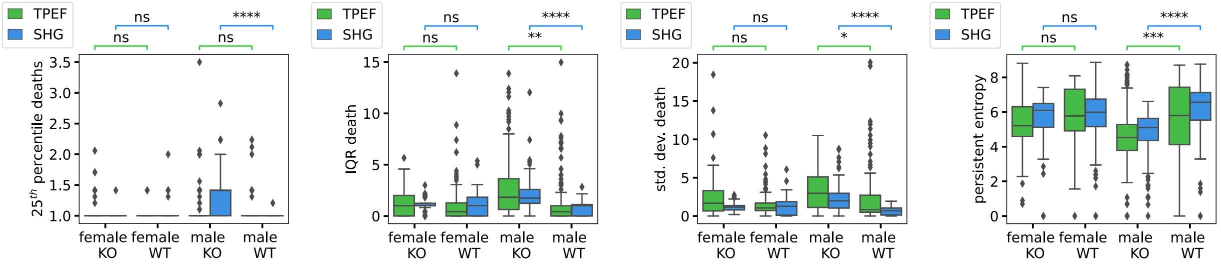


**(i)** $25^{th}$ percentile death **(j)** IQR death **(k)** std. dev. death **(l)** persistent entropy

**Figure S1.** Box plots of persistence statistics from quadrant 2 of $H_0$ per image patch, which summarise the micro-holes. The box plots show the median and quartiles, with outliers marked as points outside of 1.5 times the interquartile range. We annotate their significance using the adjusted $p$-values for four comparisons (male OcnVEGFKO v. male WT, female OcnVEGFKO v. female WT, male OcnVEGFKO v. male WT, female OcnVEGFKO v. female WT) on each imaging type (TPaF or SHG). Here the stars indicate significance levels: 'ns' is not significant $0.05 < p \leq 1$, '*' $0.01 < p \leq 0.05$, '**' $0.001 < p \leq 0.01$, '***' $0.0001 < p \leq 0.001$, and '****' $p \leq 0.0001$. We include only four of eight significance annotations for readability. Std. dev. is the standard deviation and IQR is the interquartile range.

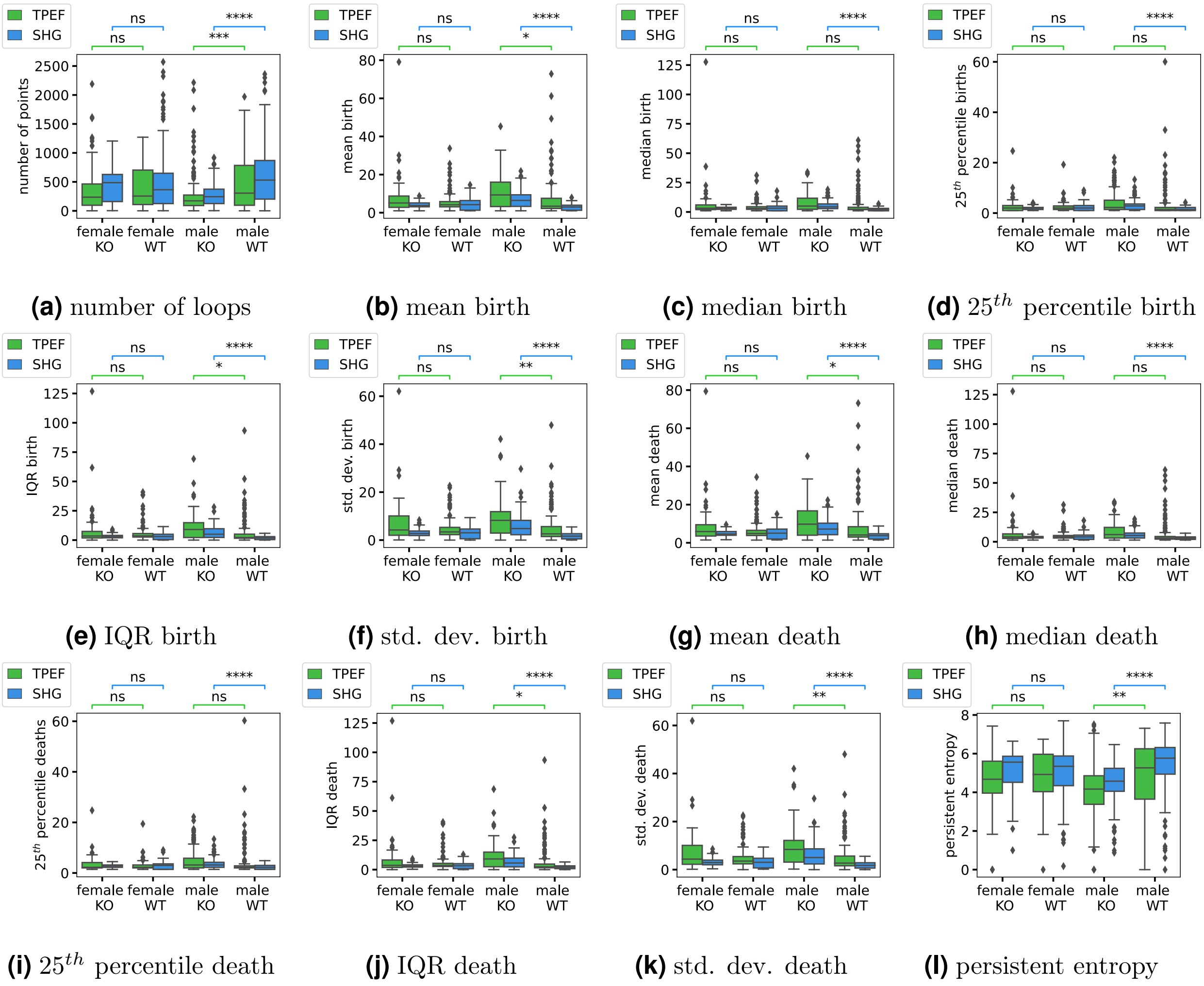


**(a)** number of loops **(b)** mean birth **(c)** median birth **(d)** $25^{th}$ percentile birth

**(e)** IQR birth **(f)** std. dev. birth **(g)** mean death **(h)** median death

**(i)** $25^{th}$ percentile death **(j)** IQR death **(k)** std. dev. death **(l)** persistent entropy

**Figure S2.** Box plots of persistence statistics from quadrant 1 of $H_1$ per image patch, which summarise the loops in the filtration that are regions of bone surrounded by micro-holes. The box plots show the median and quartiles, with outliers marked as points outside of 1.5 times the interquartile range. We annotate their significance using the adjusted $p$-values for four comparisons (male OcnVEGFKO v. male WT, female OcnVEGFKO v. female WT, male OcnVEGFKO v. male WT, female OcnVEGFKO v. female WT) on each imaging type (TPaF or SHG). Here the stars indicate significance levels: 'ns' is not significant $0.05 < p \leq 1$, '*' $0.01 < p \leq 0.05$, '**' $0.001 < p \leq 0.01$, '***' $0.0001 < p \leq 0.001$, and '****' $p \leq 0.0001$. We include only four of eight significance annotations for readability. Std. dev. is the standard deviation and IQR is the interquartile range.
